# Imaging action potential in single mammalian neurons by tracking the accompanying sub-nanometer mechanical motion

**DOI:** 10.1101/168054

**Authors:** Yunze Yang, Xian-Wei Liu, Hui Yu, Yan Guan, Shaopeng Wang, Nongjian Tao

## Abstract

Action potentials in neurons have been studied traditionally by the patch clamp and more recently by the fluorescence detection methods. Here we describe a label-free optical imaging method that can measure mechanical motion in single cells with sub-nanometer detection limit and sub-millisecond temporal resolution. Using the method, we have observed sub-nanometer mechanical motion accompanying the action potential in single mammalian neurons. The shape and width of the transient displacement are similar to those of the electrically recorded action potential, but the amplitude varies from neuron to neuron, and from one region of a neuron to another, ranging from 0.2 - 0.4 nm. The work indicates that action potentials may be studied non-invasively in single mammalian neurons by label-free imaging of the accompanying subnanometer mechanical motion.

## Introduction

Action potentials (APs) play a central role in neuronal communication^1^. While most studies to date have been focused on recording AP as an electrical signal, theories have predicted that linear and non-linear (e.g., solitons) mechanical waves accompany AP propagation along the axons^2, 3^, and the interplay between the electrical and mechanical responses is relevant to physiological functions of the neurons^4–7^. Experimental evidence of the mechanical responses has been found in animal nerve bundles and cultured invertebrate neurons^8–15^, but measuring such mechanical responses in single mammalian neurons is challenging. To study the transient, local and subtle mechanical responses in single mammalian neurons, the measurement method must have high temporal (milliseconds) and spatial (microns) resolutions, and extremely low detection limit. Atomic Force Microscope (AFM)^16–19^, piezoelectric nanoribbons^20^ and various optical approaches^21–26^ have been applied to measure cellular mechanical deformation, but imaging the transient mechanical responses associated with AP in single mammalian neurons has not been reported.

Here we report on imaging and detection of mechanical motion that accompany the AP in single mammalian neurons using an optical method that can resolve mechanical displacement with sub-millisecond temporal resolution and sub-nanometer detection limit. We study neuron-to-neuron variability, and local variability within a neuron in both the amplitude and direction of the membrane displacement. We compare the results with the simultaneously recorded action potentials and theoretical models. We demonstrate the method for non-invasive imaging of the electrical activities in mammalian neurons by tracking the subnanometer motion.

## Results

We cultured rat hippocampal neurons on a poly-D-lysine coated glass coverslip placed on the objective of an inverted optical microscope (Fig. 1a). To initiate the AP, we patched the neuron with the standard patch clamp technique, recorded the electrical signal and imaged the associated mechanical displacement of the neuron simultaneously with a differential optical detection algorithm described below (Fig. 1b) ^27^. The optical image reveals individual neurons, and zooming-in of a neuron shows the edge of the neuron. We selected a rectangular region of interest (ROI) that includes the edge, and divided the ROI along the edge into two equal halves: one half is inside of the neuron, and the second half falls outside of the neuron (Fig. 1b). We denoted the intensities of the two halves as I_1_ and I_2_, respectively. If the neuron expands, then I_1_ decreases and I_2_ increases. Conversely, if the neuron shrinks, then I_1_ increases and I_2_ decreases. The membrane displacement is proportional to (I_1_−I_2_)/(I_1_+I_2_) within a displacement range of several hundred nm (Supplementary Information, Fig. S1). We determined the range, and calibrated the relation between the displacement and normalized differential intensity, (I_1_−I_2_)/(I_1_+I_2_), using the procedure described in Supplementary Information.

**Figure 1:**
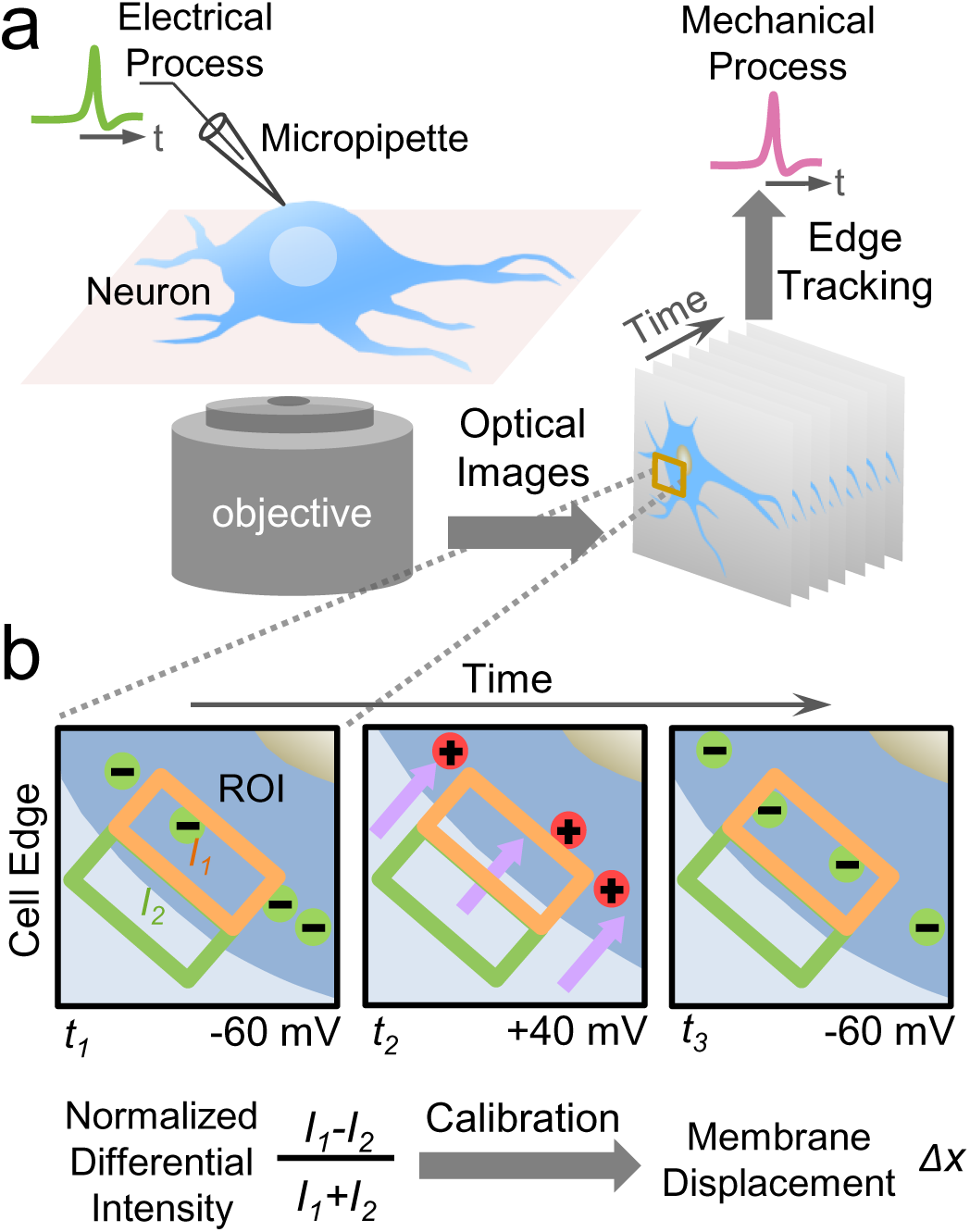
Optical imaging of mechanical motion accompanying action potential in single mammalian neurons. a) Experimental setup showing a hippocampal neuron cultured on a glass slide mounted on an inverted optical microscope, which is studied simultaneously with the patch clamp configuration for electrical recording of the action potential, and optical imaging of membrane displacement associated with the action potential. b) Imaging and quantification of membrane displacement with a differential detection algorithm that tracks the edge movement of a neuron. A region of interest is selected to include a portion of the neuron edge, which is divided into two halves along the edge (orange and green boxes) with intensities, I_1_ and I_2_, respectively. The expansion or shrinkage of the neuron is measured by a membrane displacement, Δx, from the change in the intensities, which is proportional to the normalized differential intensity, (I_1_−I_2_)/(I_1_+I_2_). See text for details.

By measuring the normalized differential image intensity, (I_1_−I_2_)/(I_1_+I_2_), within each ROI, we obtained the local membrane displacement at every location along the edge of the neuron (Fig. 1b). The spatial resolution of the method is limited by optical diffraction, which is ∼0.4 μm in the present case (numerical aperture of the objective is 0.65). The differential detection algorithm in the present work reduces common noise, such as light intensity fluctuations, which leads to a detection limit of ∼0.2 nm of membrane displacement with a denoising scheme detailed in Methods and Supplementary Information. Compared to AFM that measures one location at a time, the present method is fast and allows multiple neurons and different regions of the same neuron to be imaged simultaneously.

### Optical imaging of membrane potential-induced membrane displacement

We first examined the method by applying it to measure membrane potential-induced mechanical displacement in human embryonic kidney 293T (HEK293T) cells. For this purpose, we modulated the membrane potential of a HEK293T cell sinusoidally in the whole-cell patch clamp configuration, and recorded the electrical current and optical images simultaneously (Fig. 2a). Fig. 2d shows the current response (middle panel, red trace) to an applied potential modulated between −60 and 40 mV with frequency of 23 Hz (top panel, blue trace). The current is due to polarization, as no AP is excited in HEK293T. This is also confirmed by an observed 90°-phase shift between the applied membrane potential and the measured current. Associated with the membrane potential modulation, we expect a membrane displacement according to the thermodynamic theories.^28, 29^ The membrane displacement is too small to be visualized directly from the bright field optical image. For example, Fig. 2b shows two images of the cell edge (marked with blue dashed line in Fig. 2a) recorded at −60 and 40 mV, respectively, which show no visible difference. However, difference image obtained by subtracting the image at −60 mV from that at 40 mV shows clear contrast of the cell edge with an orange and a purple bands in Fig. 2b, corresponding to an increase and a decrease in the image intensity, respectively (see Methods for obtaining difference image). This distinct contrast in the difference image reflects a membrane displacement, which can be understood by plotting intensity profiles across the cell edge at −60 (solid orange line) and 40 mV (dashed orange line). The difference between the two intensity profiles is a lateral displacement (Fig. 2b). Subtraction of the two intensity profiles leads to regions with increased or decreased intensity, which gives rise to the orange and purple bands in the difference image.

**Figure 2:**
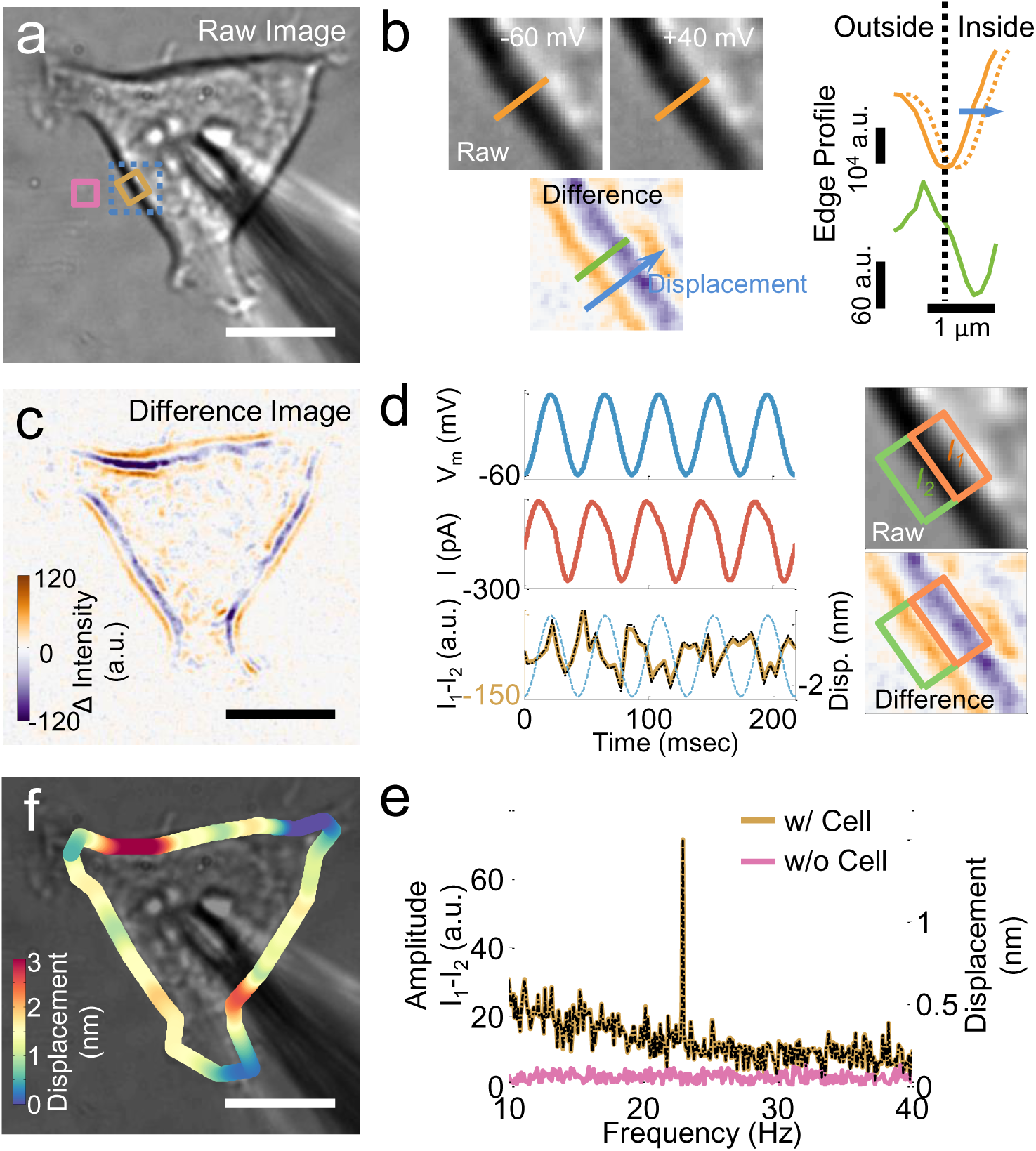
Relationship between membrane potential and cellular membrane displacement. a) Bright field image of a HEK293T cell and a micropipette used to change the membrane potential. b) Left panel: Zooming-in of the bright field images (grey scale) of the cell edge at −60 and 40 mV, and difference image (colored) obtained by subtracting the bright field image obtained at −60 mV from that at 40 mV. The characteristic orange and purple lines represent the cell edge displacement. Right panel: Corresponding intensity profiles along lines indicated in the images, where the solid and dashed orange lines are from the bright field images at −60 mV and 40 mV, and the solid green line is from the difference image. c) Difference image of the entire cell showing cell edge displacement and local variability (intensities for the orange and purple bands) of the displacement. d) Left panel: Time traces of applied potential at frequency 23 Hz (blue), measured current (red) from the patch clamp setup, and differential intensity (I_1_−I_2_, brown) from the optical imaging method. Note that the actual displacement determined from the differential detection algorithm is shown as a black line overlaid on top of the differential intensity, and the potential is indicated as dashed blue line. Right panel: A region of cell edge shown in raw bright field (grey scale) and difference (colored) images. e) Fast Fourier Transformation (FFT) power spectra of differential intensity in regions with (brown) and without (pink) cell (as indicated as colored squares in a). The FFT power spectrum of the membrane displacement is shown as a black dashed line. f) Membrane displacement map obtained from the differential detection algorithm. **Scale bar** in a, c, f): 10 μm.

The analysis above demonstrates that the potential-induced membrane displacement can be visualized by the difference image. Fig. 2c is the difference image of the cell shown in Fig. 2a, revealing the contrast as the characteristic purple and orange bands along the cell edge. According to the differential detection algorithm, the intensities of the purple and orange bands are I_1_ and −I_2_, which provide membrane displacement. The upper right of Fig. 2d shows a ROI at a cell edge, with a half inside (I_1_) and another half outside (I_2_) of the cell, and the lower right of Fig. 2d is the difference image of the same region. I_1_−I_2_ is the sum of the purple and orange bands in the difference image, which is proportional to the membrane displacement. Fig. 2d (lower left panel) plots I_1_−I_2_ vs. time, showing a periodic oscillation due to membrane depolarization-induced cell edge movement. Fast Fourier Transform (FFT) of the I_1_−I_2_ trace reveals a pronounced peak at 23 Hz, the frequency of potential modulation (Fig. 2e). The peak amplitude in the FFT spectrum provides an accurate measurement of the membrane displacement. The FFT analysis also reveals a 180**°** phase shift between the membrane potential and displacement, indicating that the cell edge shrinks as the membrane depolarizes.

We validated the above results by carrying out further control experiments and analysis. First, we did not detect any peak at the frequency of modulation in the FFT spectrum in regions outside the cell (pink line, Fig. 2e), indicating that the image intensity change at 23 Hz was due to the mechanical response of the cell to the membrane potential modulation. Second, we did not detect any peak in the FFT spectrum in the cell region when we turned off the membrane potential modulation (Fig. S3), demonstrating that the observed membrane displacement was truly originated from the membrane potential modulation. Finally, we detected little signals at frequencies other than the modulation frequency in the FFT spectrum, which shows that the measured potential-induced membrane displacement has sufficient signal-to-noise ratio compared to the background micro-motions of the cell arising from thermal fluctuations and metabolic activities (Fig. S2).

### Quantification of membrane potential-induced mechanical response

The analysis above shows that subtle cell membrane displacement can be visualized in the difference image, and measured by tracking the image intensity change along the cell edge with the differential detection algorithm. To quantify the displacement, we performed calibration using the procedure described in Methods and Supplementary Information (Fig. S1). The calibration allowed us to determine the local membrane displacement from the differential image intensity, (I_1_ − I_2_)/(I_1_ + I_2_). Scale bars in Figs. 2d and 2e show the membrane displacement of the region shown in Fig. 2a. The membrane displacement is about 1.5 nm, corresponding to 15 pm per mV. The signal to noise ratio (SNR) is ∼7, as determined from the peak amplitude at the modulation frequency, and the background level near the modulation frequency, which leads to a noise level of ∼0.2 nm of the region. The averaged displacement along the edge of the cell is ∼1.4 nm. This number is consistent with model estimate based on the Lippmann equation ^17^.

Because different regions of a cell have different membrane displacements, we plotted the local membrane displacement of a cell in Fig. 2f, showing large variability in membrane displacement along the cell edge. Some regions displace by as many as several nm (red), while other regions by only a fraction of nm (dark blue). Heterogeneity in cellular membrane is well documented in literature, including expression level of membrane proteins, type of lipids and structure of cytoskeleton, which play various functional roles in the cell ^30^. The present study reveals large heterogeneity in the membrane potential-induced mechanical response^31^.

### Imaging of mechanical motion accompanying action potential

We have demonstrated that potential-induced membrane displacement in cells can be imaged with sub-micron spatial resolution, and quantified with sub-nm precision. We now turn to the detection of mechanical motion associated with the AP propagation in primary neurons. We first identified a neuron with the bright-field optical microscope, and patched it with the patch clamp setup (Fig. 3a). We then triggered the AP by injecting current into the neuron with the patch clamp micropipette, and recorded both the electrical potential (Fig. 3b), and a stack of optical images with a frame rate of 1603 fps. To visualize the mechanical response, we obtained difference images by subtracting the first image from each of the images in the stack (see Methods). The difference images can be viewed as a movie (Supplementary Information), which shows the transient mechanical response of the neuron associated with AP. From the video, we can visualize the AP initiation and progression in the regions of axons and dendrites. This spatial information is not previously accessible by the traditional electrical recording or other methods.

**Figure 3:**
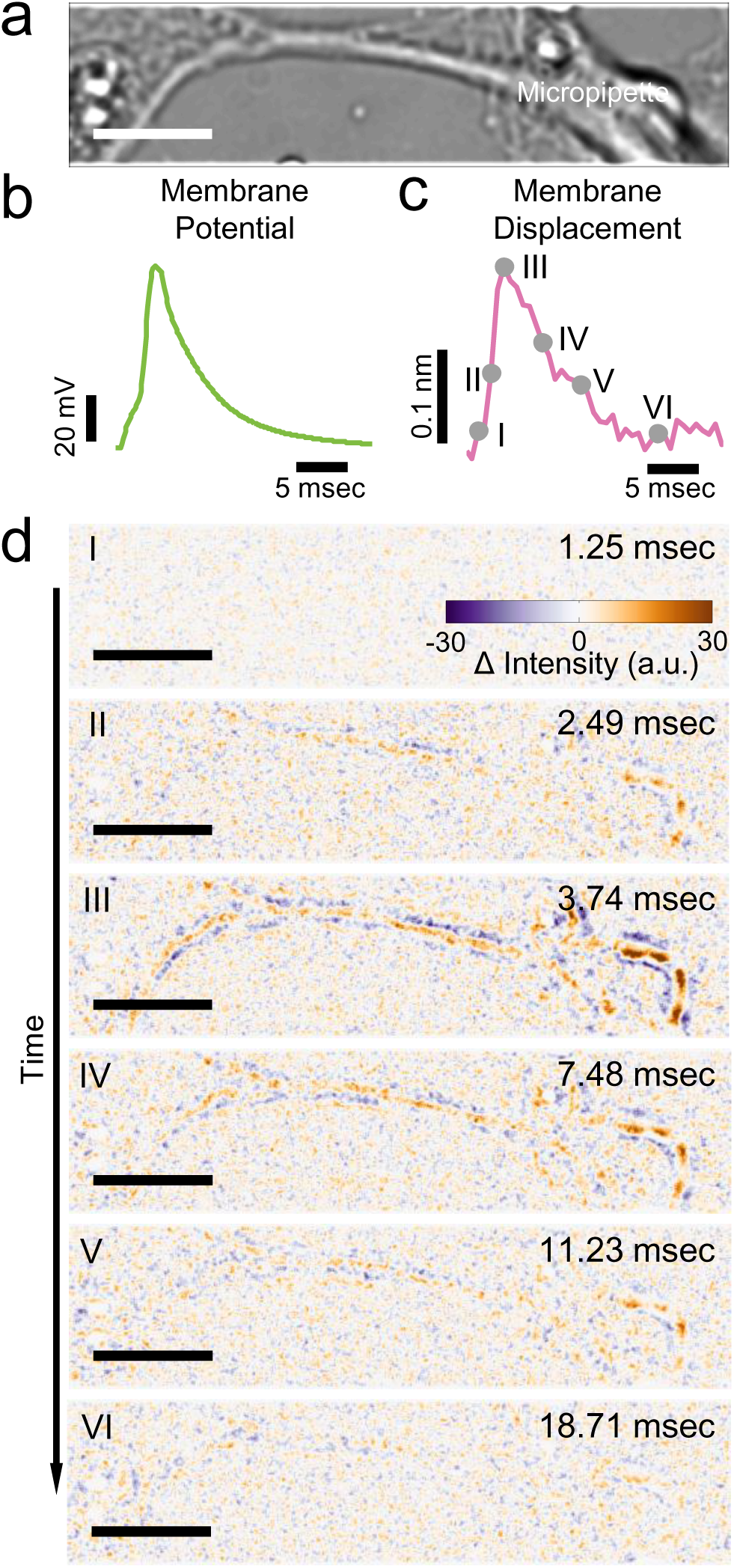
Mechanical displacement accompanying action potential in single mammalian neurons. **a)** Bright field image of a hippocampal neuron. **b)** Action potential of the neuron (green). **c)** Simultaneously recorded membrane displacement of the same neuron (magenta) with 440 traces averaged. **d)** Snapshots of difference images of the hippocampal neuron at different stages of the action potential spike. Filled grey circles in c) indicate the moments where the difference images were recorded. **Scale bar:** 10 μm.

Fig. 3d shows a sequence of difference images captured at different stages of an AP (marked on Fig. 3c). Before firing of the AP, the difference image is featureless (Fig. 3d, I). As the AP rises, the difference image begins to show contrast (Fig. 3d, II), and the image contrast reaches a maximum at the peak of the AP (Fig. 3d, III). When the AP falls, the difference image contrast also decreases (Fig. 3d, IV, and V), and eventually disappears to the background noise level (Fig. 3d, VI). Note that the purple and orange bands in the difference images in Fig. 3d represent the decrease and the increase in the image intensity, which is expected for the membrane displacement (expansion or shrinking) as we discussed earlier.

We quantified the membrane displacement using the differential detection algorithm and obtained the overall transient displacement during the AP. To clearly resolve the small displacement in the presence of micro-motion of the mammalian neurons, we evoked the AP repeatedly and obtained the average transient displacement (Fig. 3c), which shows membrane displacement with waveform similar to that of the simultaneously recorded AP (see Methods and Supplementary Information for the procedure). The maximum amplitude of the membrane displacement is ∼0.2 nm (at ∼90 mV depolarization potential), taking place at the peak of the AP. A recent mechanical theory predicted an AP induced membrane displacement in neurons^2^. For hippocampal neurons, the theory estimated a maximum membrane displacement of ∼0.1 nm, close to the present observation. However, the predicted mechanical waveform contained an oscillatory feature with a small negative displacement region followed by a more dominant positive displacement region, which was in contrast to the present observation that mechanical and electrical waveforms were similar.

The observation of membrane displacement with amplitude of ∼0.2 nm shown in Fig. 3c is an averaged value over the entire neuron. The optical method here also provides valuable local information, which reveals large variability in the local membrane displacement, not only in the amplitude but also the polarity. Fig. 4b shows the local displacement along the edge of a neuron (see Fig. 4a for the optical image). In some regions (e.g., region 1), the displacement is negative (blue, cell membrane moves inward), indicating shrinking of the neuron. In other regions (e.g., region 2), however, the displacement is positive (red, cell membrane moves outward), which is expected for expansion or swelling of the neuron. There are also regions (e.g., region 3) showing positive displacement on one side and negative displacement on the opposite side, suggesting a displacement of the entire section of the neuron in one direction. Fig. 4c plots the membrane displacement waveforms at different locations. Despite the polarity difference, the waveforms are similar in shape to each other, and also to the electrically recorded AP.

**Figure 4:**
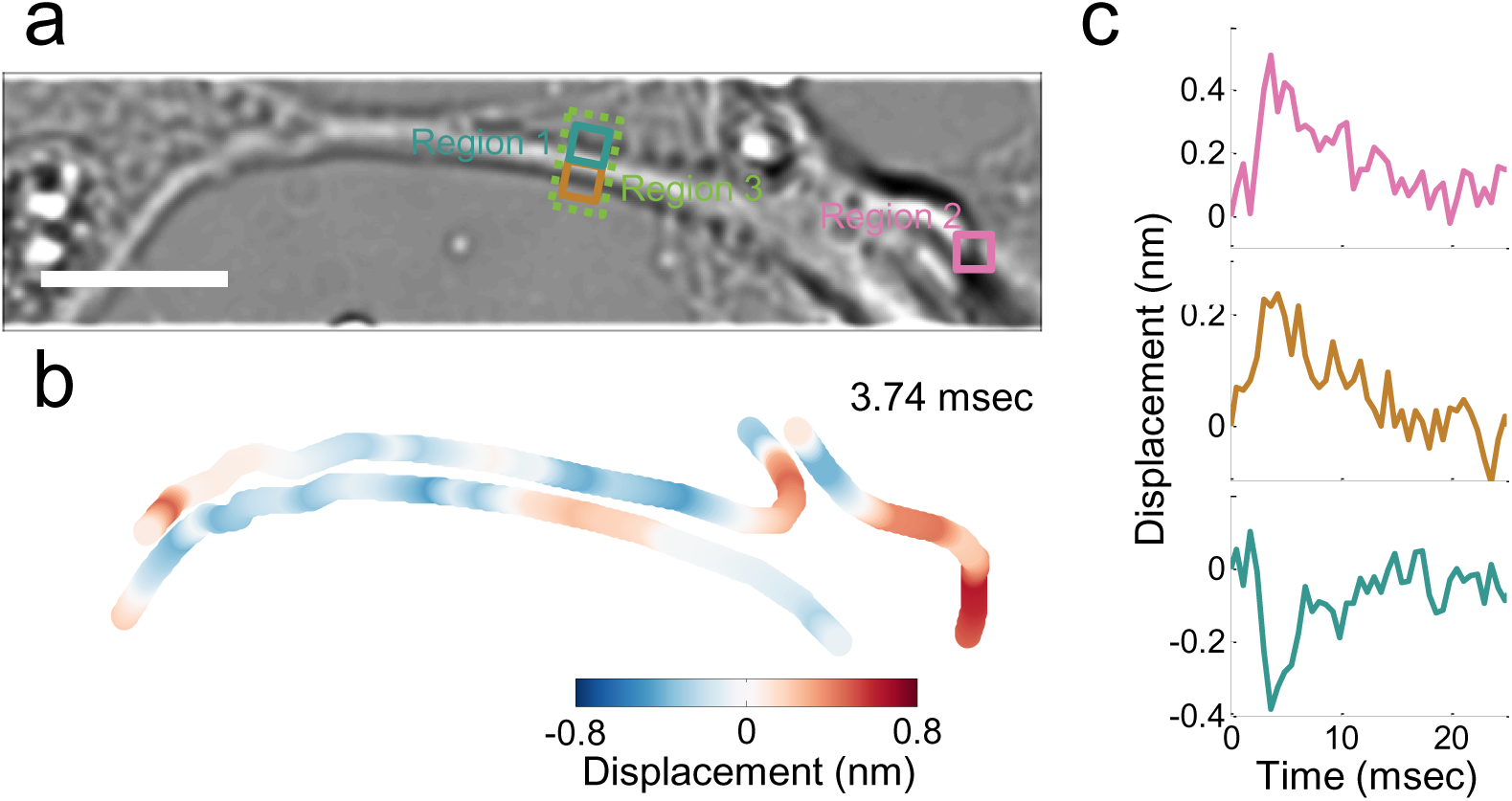
Mapping and quantification of local membrane displacement accompanying an action potential. **a)** Bright field image of a hippocampal neuron. **b)** Local membrane displacement map at the peak of the action potential, where red indicates outward displacement, blue indicates inward displacement. **c)** Mechanical waveforms at locations indicated by the corresponding colored squares in a). **Scale bar:** 10 μm

Although the observed mechanical displacement associated with the AP is predicted by recent theories, the large local variability in the mechanical deformation calls for more detailed models. The theories assume a uniform cylinder for the neuron axon, and consider primarily elastic energy associated with membrane surface area changes^2^. Elastic energy may also be stored in the membrane curvature that is related to bending modulus^29, 32^. Additionally, a more complete theory may also need to consider the local structural and mechanical variability in neurons, cytoskeletons^33^, as well as the interaction of neurons with the supporting surface. Water molecules and ions flow in and out of the neurons could also contribute to the observed membrane displacement^11, 34^.

We imaged the mechanical deformation in 11 different neurons, and observed mechanical displacement that accompanies the AP in every neuron (Fig. 5). We also observed the mechanical displacement in cultured rat Dorsal Root Ganglion (DRG) neurons (Supplementary Information, Fig. S4), which suggests that the phenomenon is a robust. Our data also show that despite both positive and negative membrane displacements are observed within a neuron, the absolute displacement averaged over a whole neuron displays a waveform that matches well with the simultaneously recorded AP. The average maximum displacement is ∼0.3 nm, corresponding to ∼4 pm/mV, but the maximum displacement amplitude varies from ∼0.2 to ∼0.4 nm, from neuron to neuron (Fig. 5). The width of the mechanical waveform matches that of the AP for each neuron (Fig. 5).

**Figure 5:**
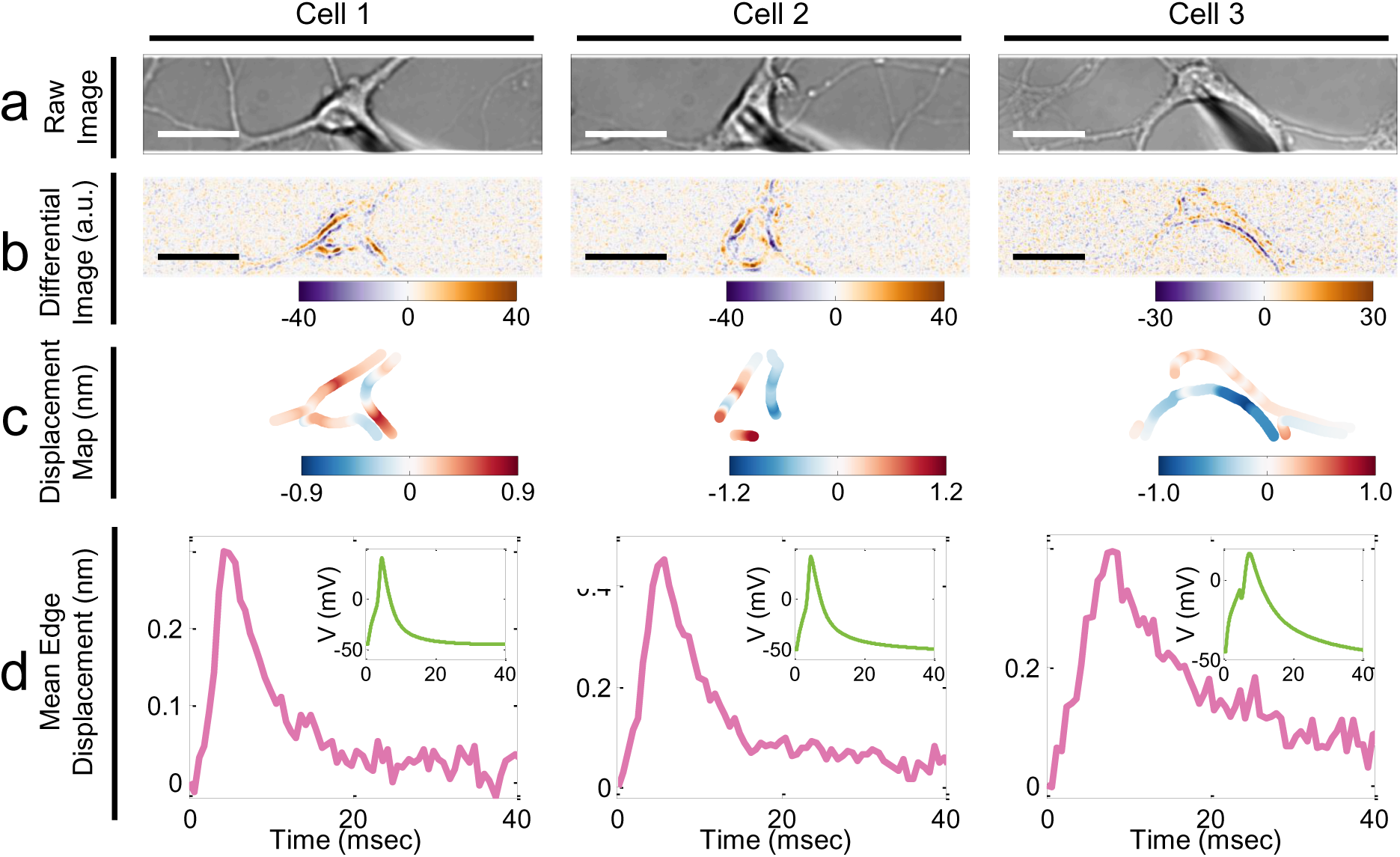
Variability of mechanical displacement in different neurons. Bright field images (a), difference images (b), local membrane displacement maps (c), and mean membrane displacement profiles (d) of three hippocampal neurons during action potential firing. The simultaneously recorded corresponding action potential profiles for the neurons are presented in the insets of (d). **Scale bars:** Cell 1 and Cell 2: 15 μm; Cell 3: 20 μm.

The data discussed above show that the AP is associated with a mechanical motion in the cell membrane, and the corresponding membrane displacement is a fraction of nm. To optimize the detection of such a small displacement, we analyzed different noise sources. At high frequencies (>200 Hz), shot noise dominates (Supplementary Information, Fig. S5), which can be reduced by increasing pixel size (or number) and light intensity. However, at low frequencies (<200 Hz), the intrinsic cell micro-motions dominate the noise spectrum as shown in the Supplementary Information (Fig. S6 and Fig. S7).

## Conclusions

We have imaged local membrane displacement at sub-nanometer level in single mammalian neurons during the initiation and propagation of the action potential. The mechanical displacement and AP have similar shapes and widths, and their peaks are synchronized with each other within the temporal resolution (0.6 ms) of the present imaging method. The mechanical displacement is observed in different neurons, indicating the phenomenon is robust, but the maximum displacement amplitude in different neurons are different, varying between 0.2 - 0.4 nm. There is also large variability in the local membrane displacement along the edge of a neuron, in both amplitude and direction: some regions expand while others shrink. Despite the local variability, the absolute displacement of a neuron displays a waveform similar to that of the action potential. While mechanical waves are anticipated to accompany action potentials by theories, detailed descriptions of the waveform, and the local variability in the membrane displacement call for more detailed theories. Since mechanical motions are always coupled with electrical signals in neurons^35^, our work demonstrates a label-free approach to image the electrical activities in neurons by tracking the mechanical responses.

## Methods

### Optical imaging of membrane displacement

The membrane displacement was measured with an inverted optical microscope (IX81) using a 60x objective (NA = 0.65), a halogen lamp in the Köhler configuration, and an sCMOS camera (Hamamatsu, Flash 4.0) for imaging. To image the membrane displacement in HEK293T cells, the camera was operated with a frame rate of 400 fps and pixel density of 256 x 512 (2 pixel binning mode). For imaging action potential-induced mechanical motions in neurons, the frame rate of the camera was increased to 1603.5 fps with pixel density of 128 x 512. Images were streamed to a fast disk array consist of 4 solid-state hard disks (Samsung 840 Pro 512GB) in parallel writing mode (RAID 0). To minimize the shot noise, the image intensity captured by the camera was adjusted close to camera saturation. To synchronize the electrical recording with images, the camera was externally triggered by the patch clamp system. The whole setup was placed on a floating optical table (Newport RS2000) and enclosed in a custom built faradic cage and acoustic enclosure to minimize electrical and mechanical noises.

### Electrophysiological recording

Electrophysiological signals were recorded in the whole-cell configuration using Axopatch 200B amplifier (Axon Instruments). For imaging membrane displacement in HEK293T cells, membrane potential was modulated sinusoidally in the voltage-clamp mode with typical frequency of 23 Hz for 10 seconds (otherwise indicated in manuscript). For action potential imaging in hippocampal neurons, multiple depolarization current pulses (typically 500 pA, 2–4 msec) were injected into the cell to evoke action potential trains at 23 Hz for 20 seconds (otherwise indicated in manuscript). Glass micropipettes were prepared with a pipet puller (P-97, Sutter Instrument) with a typical resistance of 3–8 MΩ. Before patching a cell, the micropipette was filled with intracellular recording solution consisting of 10 mM NaCl, 135 mM K-gluconate, 10 mM HEPES, 2 mM MgCl_2_, 2 mM Mg-ATP and 1 mM EGTA (pH 7.4). Cells and neurons were recorded in extracellular recording solution, containing 135 mM NaCl, 5 mM KCl, 1.2 mM MgCl_2_, 5 mM HEPES, 2.5 mM CaCl_2_ and 10 mM glucose at pH 7.4. The electrical signals were filtered by a 1 kHz low-pass Bessel filter.

### Cell and neuron culture

Wildtype HEK293T cells (ATCC, CRL-3216) were seeded on a fibronectin (Sigma-Aldrich, f1141) coated petri dish one day before the experiment. Cells were cultured in DMEM (Lonza) medium with 10% FBS (Life Technologies, 10437077) according to the user’s manual from ATCC. E18 rat hippocampus (Brainbits) were dissociated with papain and seeded on a poly-D-lysine (Sigma-Aldrich, P6407) coated glass coverslip at a density of ∼10^5^ neurons/cm^2^. After incubation in growth medium (NbActive4, Brainbits) in 5% CO_2_ humidified atmosphere at 37°C for 5–8 days, the neurons were ready for experiment.

### Differential detection algorithm

The optical images were first interpolated five times in size by adding additional pixels. The image size represented by each pixel size was determined by dividing the physical dimension of the pixel in the sCMOS imager (6.5 μm, Hamamatsu Flash 4.0) with the optical zoom and digital interpolation in the system. An ROI with a typical size of 100 x 100 pixels (2.2 μm x 2.2 μm) was selected at one location at the cell edge and split into two halves, one inside (I_1_) and one outside (I_2_). For calibrating the response of normalized differential intensity with displacement, the ROI was shifted by a certain number of pixels in the direction perpendicular to the edge (Fig. S1). Normalized differential intensity, (I_1_−I_2_)/(I_1_+I_2_), vs. pixel shift (displacement) showed a typical “S” shape (Fig. S1). Within a certain range (marked between two dashed lines in Fig. S1d, ∼500 nm), the differential intensity change is linearly proportional to the pixel shift (displacement). The slope of the linear region was determined, and used as the calibration factor to determine the membrane displacement with the differential detection algorithm (Fig. S1e). For all the measurements performed in this work, the membrane displacement was small (less than few tens of nm) and well within the linear region (Fig. 2d).

For both HEK293T cells and hippocampal neurons, cell edges were manually identified, and the local membrane displacement was determined using the procedure described above. Note that the differential detection algorithm is insensitive to the accuracy of edge identification as long as the edge falls within the ROI, and the displacement is within the linear rage (∼500 nm) as shown in Fig. S1e. After the displacement at one location we quantified, the ROI was automatically shifted to an adjacent location, and the procedure was repeated until the displacement along the edge of the entire cell was determined. In the case of HEK293T cells, the ROI was shifted every 10 degrees to the centroid of the cell (average ∼1.7 μm/step along the edge for 10 μm cell in radius). For neurons, the ROI was shifted every 64 pixels along the boundary (∼1.4 μm/step for 60x zoom). It took about 5 mins to complete the analysis of one cell with the method, which can be further optimized in the future.

Edge tracking of cells was reported to detect membrane flickering via tracking the derivative of the intensity profile across a cell edge^36^. Rather than tracking the derivative, we determined the intensity difference between two regions. Because the intensity in each region was integrated over many pixels, our approach minimized pixel noise (shot noise). We further normalized the intensity difference (I_1_−I_2_) by the total intensity (I_1_+I_2_) to remove the common noise from the light source, allowing imaging of the sub-nanometer membrane displacement in neurons.

### Denoising scheme and difference image

To resolve the subtle mechanical displacement, denoising procedures were applied to measurements of both HEK293T cells and hippocampal neurons. In the HEK293T experiment, because a sinusoidal potential was used to modulate the membrane, Fast Fourier Transform (FFT) filter was used to remove noise at frequencies other than the modulation frequency for each ROI (Supplementary Information). The amplitude and phase at the modulation frequency were quantified as magnitude and direction of the edge displacement while membrane potential depolarized.

To obtain the time difference image of the cell, FFT filter was applied to the image stack over time pixel by pixel. Since the mechanical response was in phase (0 or 180 degrees phase shift) with the applied potential modulation, real part image at the modulation frequency was selected as the time difference image (the imaginary part measured response in +/−90 degrees). Images were spatially smoothed with 3×3 pixels (0.33×0.33 μm) mean kernel before FFT filtering. Mathematically, the time difference image (quantify the temporal information first) is equivalent to the differential detection scheme (quantify the spatial information first, then denoising along temporal dimension), but it provides a convenient way to visualize the membrane displacement (both direction and amplitude).

For hippocampal neurons, repeated APs were evoked with typically 100–400 cycles. Both electrical signal and images were averaged over the repeating cycles, and an averaged signal was obtained. To observe the mechanical displacement, time difference images were obtained by subtracting the first image from the averaged image stack. Mechanical displacement was quantified using the differential detection algorithm along the edge as described earlier from the averaged image stack. To remove spatial noise, the mean image stack was temporally detrended pixel-by-pixel using a fitted linear function, and digitally filtered with a FIR low-pass filter with a cut-off frequency of 500 Hz.

### Displacement quantification and data analysis

Local membrane displacement map was obtained by interpolating local displacement quantified by the differential detection algorithm. The displacement power spectral density and standard deviation between 20 - 200 Hz were quantified as the noise level. All post-acquisition analysis was carried out by custom-written MATLAB (Natick, MA, USA) code.

## Acknowledgements

We thank Drs. Ming Gao and Jie Wu from Barrow Neurological Institute for technical support on electrophysiological recording. We thank NIH (1R01GM107165), Gordon, Betty Moore Foundation, and NSFC (Grant No. 21327008, 21327902, and 21676260) for financial support.

## Author Contributions

Y.Y. and X-W.L. designed and conducted the experiments. Y.Y. analyzed the data. N.T., Y.Y., X-W. L. and S.W. discussed results. Y.Y. and Y.G. wrote Matlab scripts. N.T., Y.Y., and X-W.L. wrote the manuscript. N.T. conceived and supervised the project.

## Additional Information

**Competing financial interests:** The authors declare no competing financial interests.

